# Structural variations in the phytoene synthase 1 gene affect carotenoid accumulation in tomato fruits and result in bicolor and yellow phenotypes

**DOI:** 10.1101/2022.10.29.514357

**Authors:** Blandine Bulot, Sébastien Isabelle, Roberto Montoya, Louis Félix Nadeau, Jonathan Tremblay, Charles Goulet

## Abstract

Tomato fruits normally accumulate large amounts of the red pigment lycopene in their chromoplasts. Some tomato cultivars (*Solanum lycopersicum*) show however distinct phenotypes, from a pure yellow hue to bicolor fruits with red and yellow sections. In this study, we show that alleles of the phytoene synthase 1 gene (PSY1), the first gene of carotenoid synthesis pathway, are responsible for the yellow, but also the bicolor phenotype. Introgression lines with the *PSY1* allele from the green-fruited species *S. habrochaites* express less the enzyme, resulting in a bicolor phenotype. On the other hand, in tomato bicolor cultivars, the same coloration pattern is caused by a 3789 bp-deletion in the promoter region of *PSY1*. Since the deletion contains part of the 5’UTR region of *PSY1*, translation efficiency is likely decreased resulting in a reduction of lycopene accumulation. Furthermore, we identified that the yellow *r^y^* phenotype is caused by a duplication and an inversion implicating *PSY1* and the downstream neighbor gene. The genomic rearrangement change the end of PSY1 amino acid sequence. The fruits of yellow *r^y^* cultivars are still able in certain conditions to accumulate lycopene near the blossom-end of the fruit, though to a lesser extent than in bicolor cultivars. In contrast, fruits of the yellow *r* cultivars never present fleshy red sections. These cultivars have an insertion of a single long terminal repeat from the Rider transposon in the first exon of PSY1 resulting in a non-functional protein. These results demonstrate how multiple phenotypes can arise from structural variations in a key gene.

## Introduction

Carotenoids, a very diversified and important group of pigments, are naturally synthesized by plants, algae, fungi and bacteria (Bartley *et al*. 1994). In plants, carotenoids can be found in chloroplasts, where they play a role in photoprotection and light harvesting for photosynthesis (Young 1991, Horton and Ruban 2004). Carotenoids also accumulate in chromoplasts, in specialized structures like plastoglobules, crystals, tubules, and inner membrane, where they are responsible of the color, from yellow to red, displayed by many flowers and fruits (Klee and Giovannoni 2011, Ljubesić *et al*. 1991). These bright colors help plants attracting pollinators and seed dispersers (Dicke and Baldwin 2010). Tomatoes are known to accumulate large amounts of carotenoids in their chromoplasts and more specifically lycopene which confers to the fruits their red color (Li and Yuan 2013).

Carotenoids are also the precursors of apocarotenoids like the phytohormones abscisic acid and strigolactones, or the volatiles safranal and β-ionone (North *et al*. 2007, López-Ráez *et al*. 2008, Ohmiya 2009). These molecules derive mostly from the oxidative cleavage of carotenoids by CCDs enzymes (Carotenoid Cleavage Dioxygenase). For example, CCDs can use lycopene as a substrate to produce the volatile 6-methyl-5-hepten-2-one (MHO) with a citrus-like aroma. The same enzymes can cleave ζ-carotene to form geranylacetone, a volatile with a fruity and fresh aroma. Apocarotenoid volatiles are known to have a positive impact on tomato flavor and increase the sweetness perception (Vogel *et al*. 2010, Tieman *et al*. 2012, Bartoshuk and Klee 2013). In addition to the apocarotenoids, about 30 volatile compounds derived from fatty acids and amino acids are thought to be essential to the distinct flavour of tomatoes (Ohmiya 2009, Ilg *et al*. 2014).

The significant color change of tomatoes during ripening, as well as the availability of a large population of colored mutants make the species particularly valuable for research on the carotenoid production pathway, which has been extensively studied. Carotenoids synthesis starts with the condensation of two molecules of geranyl geranyl diphosphate by the enzyme phytoene synthase (PSY). This reaction generates the first carotenoid, 15-cis-phytoene which is also colorless (Bartley *et al*. 1992). In tomatoes, three isoforms of phytoene synthase have been identified with different spatio-temporal expression patterns. The isoform PSY1 is expressed in the ripening fruits, while PSY2 and PSY3 are respectively present in leaves and roots (Giorio *et al*. 2008, Cao *et al*. 2019). The phytoene synthase reaction is considered as one of the most important regulatory step of the carotenoid pathway (Fraser *et al*. 2007). The carotenoid 15-cis-phytoene is then modified by a phytoene desaturase (PDS), yielding successively 15-9’-di-cis-phytofluene and 9,15,9’-tri-cis-ζ-carotene (Pecker *et al*. 1992). Subsequently, the ζ-carotene isomerase enzyme (ZISO) converts 9,15,9’-tri-cis-ζ-carotene in 9,9’-di-cis-ζ-carotene, which goes through two other desaturation reactions, managed by a ζ-carotene desaturase (ZDS) (Hirschberg 2001, Bouvier *et al*. 2005). The resulting molecule, 7,9,7,9’-tetra-cis-lycopene (prolycopene), has its configuration transformed from cis to trans by a carotenoid isomerase (CRTISO) in order to generate the carotenoid all-trans-lycopene, also called lycopene (Isaacson *et al*. 2002). At this point, the pathway can take two directions depending on the type of rings formed. The lycopene-β-cyclase (LYCB) catalyzes one or two cyclisation to yield γ-carotene and β-carotene respectively. Alternatively, the lycopene can be modified by the lycopene-ε-cyclase (LYCE) to produce δ-carotene, which can then become α-carotene by the subsequent addition of one β-ring by the lycopene-β-cyclase (Fantini *et al*. 2013).

A few tomato cultivars produce bicolored fruits that are mostly yellow at maturity but with contrasted red flesh sections. In some cultivars, these red zones are limited to the blossom end of the fruit and only occasionally appear, while other cultivars show a more consistent coloration with red streaks that can be observed in every fruit. MacArthur and Young described the first group as yellow tomatoes showing pink colors in hot summer weather (Young and MacArthur 1947). This type of coloration was suggested to be linked to the same locus as the yellow fruits (locus *r*) and the mutation was named *r^y^* (Youg 1956). Phytoene synthase 1 (PSY1) was later on identified as the gene responsible for the yellow phenotype. The color originates from the absence of carotenoid synthesis during the ripening of the fruit (Fray and Grierson 1993). The PSY1 loss of function causes a reduction of more than 99% of lycopene content in fruit, which passes from 107,5 mg per g or fresh weight in red fruits to 0,7 mg in yellow fruits (Rêgo *et al*. 1999). The *r* mutant was described as having a shorter transcript and the other yellow mutant, *r^y^*, was lacking the last 25 amino acids that were replaced by an unrelated sequence of 23 amino acids (Fray and Grierson 1993). Even if the *r^y^* locus has been well studied, the complex genomic rearrangement that caused the mutation is not fully understood. In addition, little is known about the second group of bicolor tomatoes with a stronger phenotype. In this study, we investigate further the genetic basis for yellow and bicolored fruits in tomato using heirlooms cultivars and introgression lines derived from the wild related species *S. habrochaites*.

## Results

### Mapping of a QTL for bicolored-fruits

Fruits of the tomato related wild species *S. habrochaites* do not accumulate lycopene and stay green even at maturity (Grumet *et al*. 1981). This makes introgression lines derived from *S. habrochaites* very useful to identify QTL involved in carotenoid synthesis. Introgression lines possess small chromosomal regions of wild relatives and a phenotypic difference between an introgression line and the background parent can link to the position of genes associated with a specific phenotype. In the introgression population between *S. lycopersicum* (cultivar E-6203, accession LA4024) and *S. habrochaites* (accession LA1777) developed by Monforte and Tanksley (2000), the lines LA3922 and LA3923 produce bicolor fruits **(Fig. 1A)**. According to the map published, these two lines possess one introgression of *S. habrochaites* genome at the end of the chromosome 2 (Monforte and Tanksley 2000, Mathieu *et al*. 2009). The line LA3923 possesses a shorter *S. habrochaites* introgression segment than the line LA3922 and was selected to reduce further the region responsible for the bicolored phenotype. The heterozygous F1 plants resulting from the cross between LA4024 and LA3923 produced red tomatoes confirming that the red allele is dominant over the bicolor allele. New smaller introgression lines between LA4024 and LA3923 bicolor line were selected using InDel markers. One line showed a phenotype that was incongruous with the recessive nature of the bicolor alleles, and we quickly concluded that the phenotype was not linked to the introgression on chromosome 2. The initial map of the introgression lines was done with a relatively small number of markers for each chromosome and small-undetected introgressions likely persisted in some of the lines. Since alleles of PSY1 are responsible for the yellow phenotypes, a high resolution-melting (HRM) marker was designed in the first exon of PSY1 to detect if LA3923 is having one of those undetected introgressions on chromosome 3. The genotyping results showed that plants producing red tomatoes were either homozygous for *S. lycopersicum allele* or heterozygous, while plants with bicolor fruits were all homozygous for *S. habrochaites* allele. These results suggested that the QTL is linked to PSY1 or located near the gene on chromosome 3. New introgression lines were developed from the cross between LA4024 and LA3923 to isolate the introgression on chromosome 2 (LA3923-2) and chromosome 3 (LA3923-3). The introgression line on chromosome 3 had bicolored fruits while the introgression line on chromosome 2 had red fruits. The line LA3923-3 has a small introgression on top of chromosome 3 between position 3,894,833 and 7,730,225 bp. The locus was further reduced by developing a smaller introgression on chromosome 3 (LA3923-3a) **(Supplementary Table 1)**. RNA sequencing of the parental line LA4024 and the line LA3923-3a were used to perform a SNP density analysis. Thanks to the high level of SNPs existing between *S. lycopersicum* and *S. habrochaites*, the size of the lycopene related QTL was reduced to about 676 kbp between position 3,961,150 and 4,637,328 bp. Less than 75 genes are present on this locus and only *PSY1* seems related to the carotenoid pathway. SNP density analysis was also used to verify if other small-undetected introgressions were present on other chromosomes. No introgression other than the one on chromosome 3 was identified by the analysis. Carotenoids extraction was performed on the parental line LA4024 and introgression line LA3923-3a. As expected, there is a strong decrease in lycopene (89%) in the bicolor-fruited introgression line compared with the red-fruited background, with higher content of lycopene in the red sections of LA3923-3a compared with the yellow sections (**Fig. 1B**)

**Figure 1.**
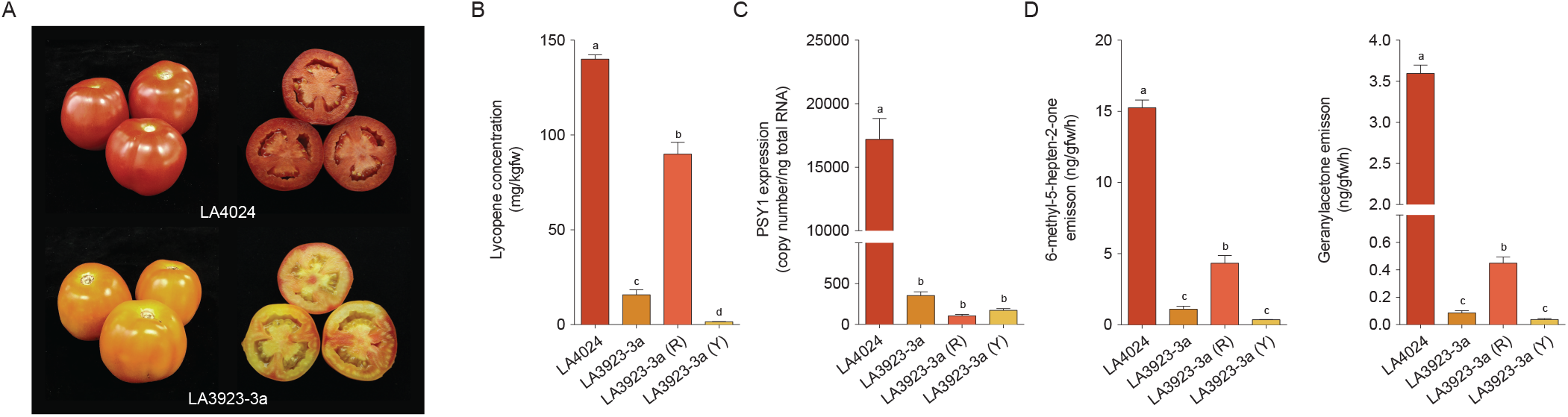
*PSY1* allele of *S. habrochaites* results in a bicolor phenotype, a reduction of lycopene, and a decrease in apocarotenoid volatiles emission. (**A**) Representative fruits of the red parental line LA4024 and the bicolor introgression line LA3923-3a. (**B**) Lycopene concentration in ripe tomato fruits of the background parent and the introgression line LA3923-3a, as well as separated yellow (Y) and red (R) sections of the bicolor line (±SE, *p*<0,05, n=4). (**C**) Transcript levels of *PSY1* in ripe tomato fruits of the parental line LA4024, LA3923-3a, and red and yellow sections of LA3923-3A. (**D**) Emission of the apocarotenoids 6-methyl-5-hepten-2-one and geranylacetone (±SE, *p*<0,05, n=6).

*PSY1* gene expression was evaluated by quantitative PCR (qPCR) on ripe fruits of LA4024 and LA3923-3a. Transcripts for PSY1 were 49-fold more abundant in the red background parent (LA4024) compared to the bicolor introgression line **(Fig. 1C).** The results were confirmed by a transcriptomic analysis (RNA-Seq) on ripe fruits from the lines LA3923-3a and LA4024. In the bicolor line LA3923-3a, the expression level of *PSY1* gene was about 2% (FPKM: 30) of the red parental line LA4024 (FPKM: 2060). These results suggest that the low expression of PSY1 is responsible for the 89% decrease in lycopene accumulation in the bicolor-fruited introgression line **(Fig. 1B)**. Yellow and red tissues from the bicolor LA3923-3a fruits were separated to estimate the expression of *PSY1* in the different sections of the tomatoes. The qPCR analysis showed no difference in *PSY1* expression between the different tissues which suggests that the reduction of *PSY1* expression happens in a global way in the fruits even if there is localized accumulation of lycopene **(Fig. 1)**. The PSY1 protein sequence from *S. habrochaites* is the same as the one from tomato except for one substitution at the end of the protein (A438V). However, in the promoter region, many deletions, insertions and SNPs were detected between the genome of two species. Those modifications are likely responsible for the change in *PSY1* expression observed in the line LA3923-3a.

To determine the impact of carotenoid accumulation on apocarotenoid volatiles emission, volatile analysis was performed on ripe fruits from LA4024 and LA3923- 3a plants. The bicolor line LA3923- 3a showed a 93% and 98% reduction in the emission of 6-methyl-5-hepten-2-one and geranylacetone respectively compared to the parental line LA4024 **(Fig. 1D)**. To further determine if there is a difference in the emission of apocarotenoids in the red and yellow tissues, bicolored fruits of LA3923-3a were cut into small red or yellow sections. Volatiles were extracted from the respective tissues. Red tissues emit significantly more apocarotenoids than the yellow tissues. More precisely, there was about 12 time more geranylacetone and 6-methyl-5-hepten-2-one in the red sections than in the yellow sections **(Fig. 1D)**.

### Structural variation in bicolor tomato cultivars

The coloration and apocarotenoid volatiles emission patterns of some bicolor heirloom and modern tomato cultivars is rather similar to what was observed in the *S. habrochaites* introgression line (**Fig. 3C, Supplementary Fig. 1**). A transcriptomic analysis was therefore performed to verify if a reduction in *PSY1* expression is also responsible for the bicolor phenotype of those cultivars. In the region identified with the introgression line, the expression of three genes was significantly different between the group of bicolor, and red cultivars; PSY1 (Solyc03g031860), an auxin responsive factor (Solyc03g031970), and a phosphoenolpyruvate carboxylase (Solyc03g032100). Among them, only *PSY1* was also less expressed in the yellow group. *PSY1* was three times more expressed in the red group than in the bicolor, yellow *r*, and *r^y^* groups (**Fig. 2**). However, in comparison, *PSY1* was 50 times more expressed in the LA4024 control line than in the bicolor introgression line LA3923-3a. Given the variation inside each group, this reduction is unlikely to explain by itself the 76% decrease in lycopene accumulation in the bicolor group (**Fig. 2B**).

**Figure 2.**
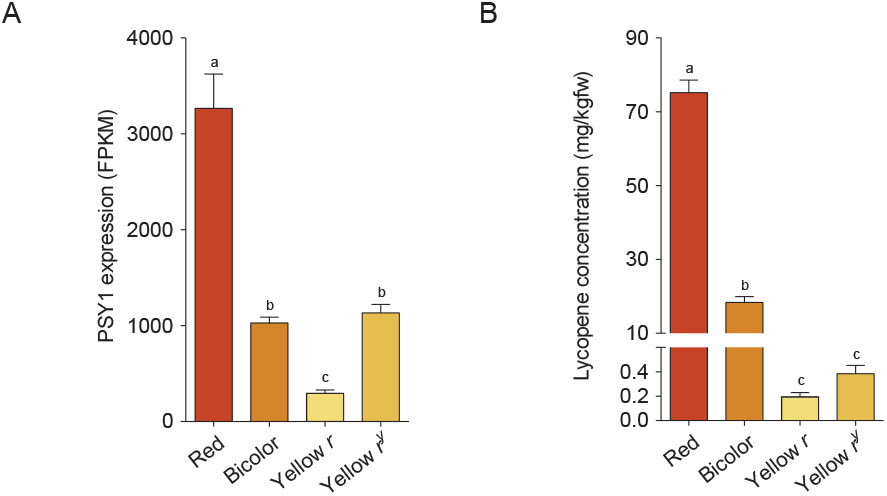
Structural variations at *PSY1* locus change the gene expression and influence lycopene accumulation in tomato cultivars. **(A)** Transcripts level of *PSY1* in ripe fruits of red, yellow (*r* and *r^y^*) and bicolor tomato cultivars. (red n=8, bicolor n=7, yellow r n=3, yellow r^y^ n=5). Significant differences between the cultivar groups are indicated with letters (±SE, *p*<0,05). **(B)**Lycopene concentration (mg/kg of fresh weight) in ripe fruits of red, yellow and bicolor tomato cultivars. The low levels of lycopene concentration in ripe fruits of bicolor and yellow cultivars cannot completely be explained by *PSY1* expression.

**Figure 3.**
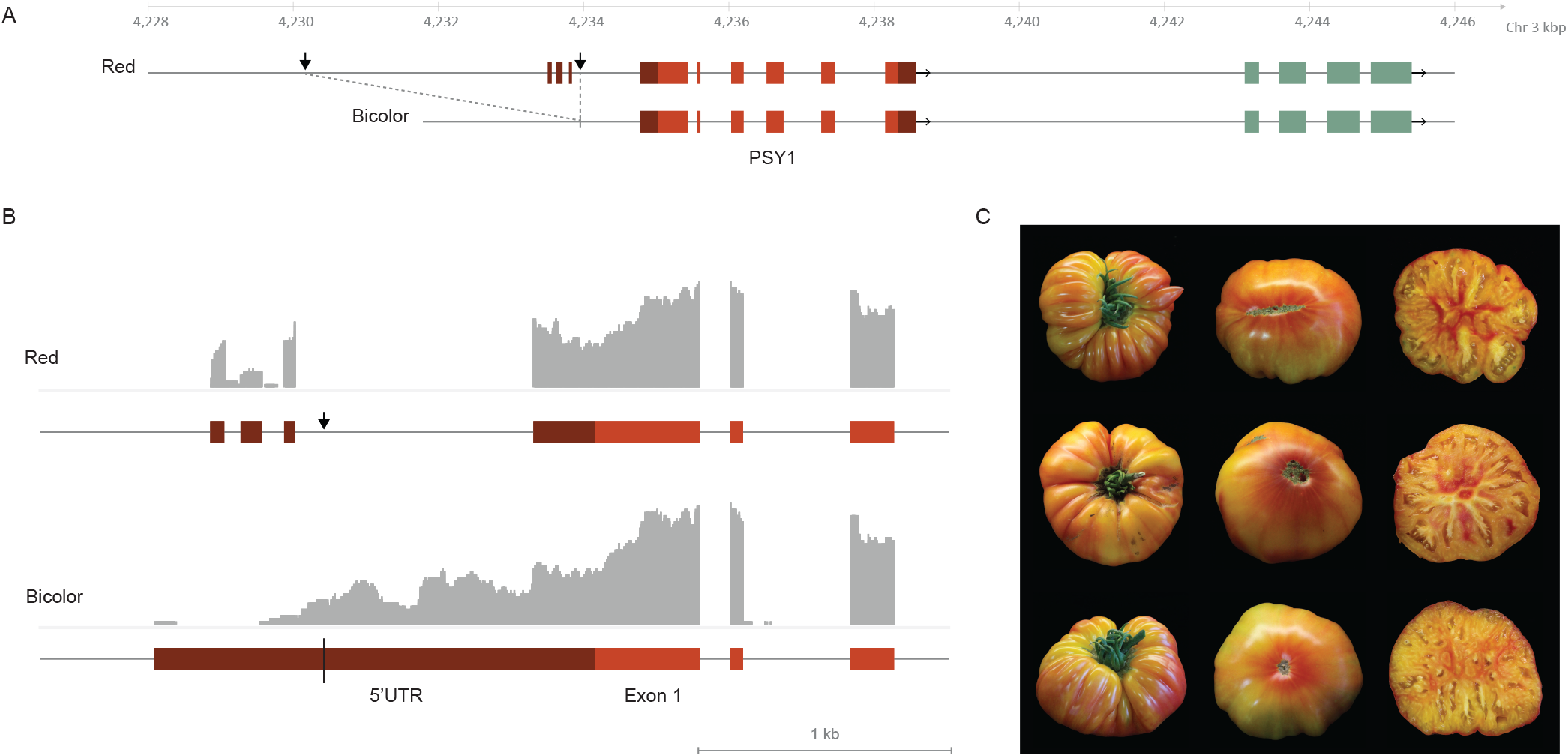
A deletion in the bicolor cultivars changes the 5’UTR of *PSY1*. **(A)** Bicolor cultivars are characterized by a 3789 bp sequence deletion upstream of *PSY1*. Exons of *PSY1* are in red, UTR are in dark red. **(B)** Transcripts alignment of *PSY1* for red and bicolor cultivars. Grey areas represent the relative read counts for each position in the transcriptomic analysis. The deletion in the bicolor cultivars results in a longer 5’UTR region with lost UTR splicing sites. **(C)** Fruit appearance of bicolor tomato cultivars. Top, bottom and cross section of the bicolor cultivars Striped German (top row), Mortage Lifter bicolor (middle row) and Grapefruit (bottom row) at ripe stage. Red tissues are present near the epidermis, in the columella, and in the septa.

Since the phenotype of yellow *r* and *r^y^* cultivars is explained by structural mutations that affect *PSY1* mRNA, an analysis of the transcripts was performed to find if variations in PSY1 sequence could also play a role in the bicolor phenotype of the cultivars. While the coding sequence of both alleles is the same, there is a major difference in the 5’ untranslated region (UTR). In the red cultivars, the 5’UTR of *PSY1* is characterized by several splicing sites with a few variants that allow to identify four 5’UTR exons (**Fig. 3B**). In the bicolor cultivars, there is a major rearrangement of the 5’UTR region caused by a deletion of 3789 nucleotides that starts 4858 bp before the start codon and finishes in the longest 5’UTR intron (**Fig. 3A**). This deletion causes a major shift in the start of the UTR. It also results in less splicing in the 5’UTR, and the 5’UTR of the bicolor cultivars is composed of only one long exon that shares only the last 244 bp with the 5’UTR of the red cultivars (**Fig. 3B**). The 5’UTR sequence of bicolor cultivars is therefore nearly 1900 bp long, while the most common splicing variant of the red cultivar, with three 5’UTR exons, is less than 350 bp. The deletion in the bicolor cultivars was further confirmed by PCR amplification of the DNA sequence.

### Genomic rearrangement in yellow tomato cultivars

*PSY1* gene sequence analysis shows that among the eight yellow cultivars studied, three have the *r* mutation (Barry Crazy Cherry, Lemon Drop and Butter Apple) while Téton de Vénus, Poma Amoris, Galina’s Yellow, Mirabelle and Gold Currant have the *r^y^* mutation. The two yellow groups, *r* and *r^y^*, show distinct versions of *PSY1* gene sequence, as well as expression and splicing variants. The cultivars of the *r* group have a 403 bp insertion in the first coding exon (**Fig. 4**). Because of this insertion, only the first 107 amino acids are conserved followed by 71 amino acids and a premature stop codon derived from the insertion. Thus, the PSY1 protein sequence of the *r* group is only 178 amino acids long, in contrast with the functional PSY1 protein that has 412 amino acids. In these yellow cultivars, PSY1 amino acid sequence is therefore severely modified and cut, which consequently leads to an inactive protein. Of the 403 bp insertion, 398 bp are part of a single long terminal repeat (LTR) normally found in Rider retrotransposons. No internal region was detected. The insertion sequence is identical to Rider LTR found in the fruit shape-related *sun* locus but has a few differences from the *r* cDNA sequence published by Fray and Grierson in 1993 (Jiang *et al*. 2009, Fray and Grierson 1993). Since all the yellow cultivars of the *r* group have the same sequence, the originally published cDNA sequence had most likely mistakes linked to the cloning or sequencing process. Transcriptomic analysis shows that *r*-yellow cultivars have a 90% reduction in *PSY1* transcripts compared to red cultivars (**Fig. 2A**). Based on the splicing counts between each exon, only a few of the transcripts going to the first exon continue to the second exon, meaning that PSY1 transcription frequently ends with the 403 bp insertion. Interestingly, a significant portion of the transcripts is directly spliced from the end of 5’UTR exon 3 to the second exon of the gene (**Fig. 4B**). This alternative splicing that removes 5’UTR exon 4 and the first coding exon was not detected in the red, bicolor and *r^y^*-yellow cultivars. The transcripts derived from this unusual splicing is unlikely to result in a functional sequence based on translation initiation predictions. In all the alternative splicing analyzed from the *r* group, none resulted in a full amino acid sequence needed for a phytoene synthase activity, explaining the lack of lycopene in the fruits of these cultivars (**Fig. 4C**).

**Figure 4.**
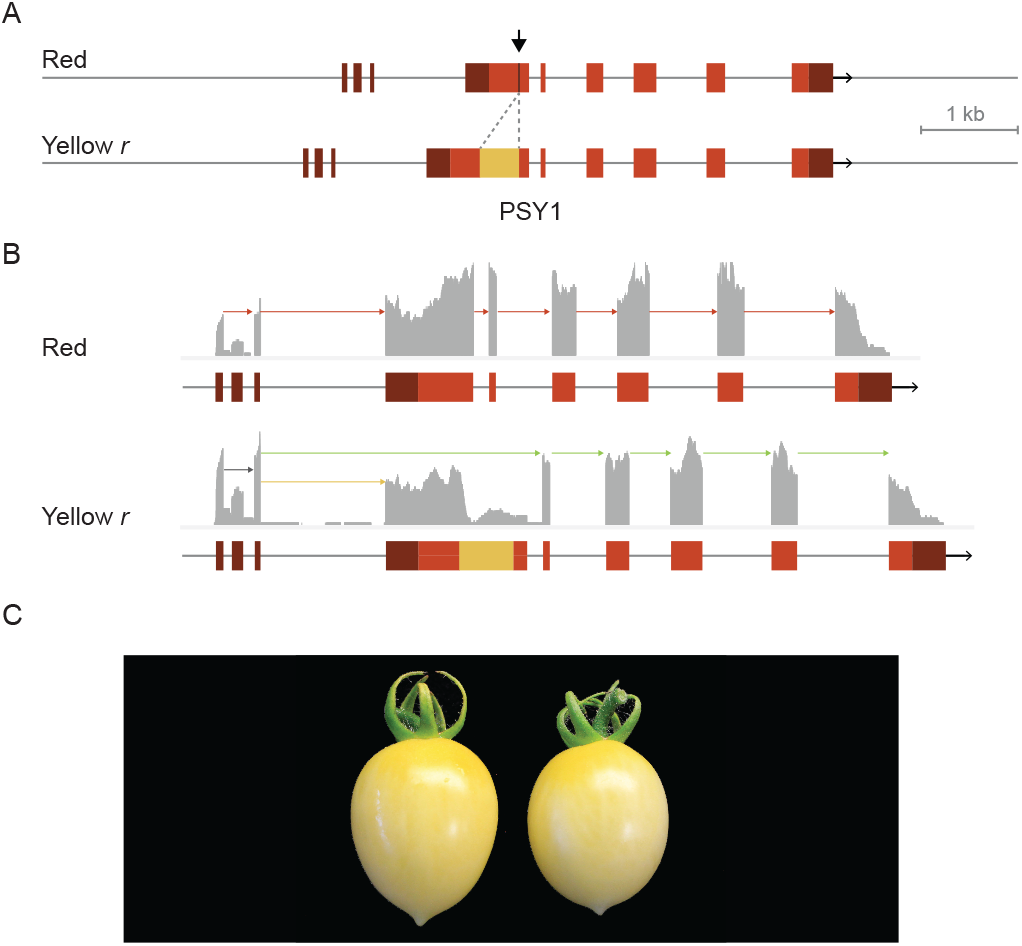
Yellow cultivars with the *r* variation do not accumulate lycopene because of an insertion in the first exon of *PSY1*. **(A)** The Rider retrotransposon insertion is indicated in yellow in the illustration. The insertion results in a premature stop codon. Exons of *PSY1* are shown in red while UTR are illustrated in dark red. (B) Reads mapping in yellow *r* and red cultivars. The insertion is linked to a decrease of read counts starting around the insertion site and only a small fraction of the transcripts continue to the second exon. An unusual alternative splicing (green arrows) was only detected in yellow *r* cultivars. The splicing links the 5’UTR directly to the second exon. **(C)** Yellow r cultivars do not visibly accumulate lycopene.

Fray and Grierson (1993) suspected that PSY1 sequence of yellow cultivars of the *r^y^* group is the result of a complex genomic rearrangement with the gene downstream of *PSY1*, an Acyl-CoA synthetase (Solyc03g031870). Using *r^y^*-yellow SRA data we investigated the genomic sequence and found that PSY mRNA sequence modification derived from an inversion and a duplication (**Fig. 5**). The rearrangement starts 98 bp before *PSY1* usual stop codon. The inversion begins with the first 191 bp of Solyc03g031870 exon 2 and continue until position -4,994 bp of *PSY1* start codon. The inverted-duplicated sequence ends in the intronic region between Solyc03g031870 exon 1 and 2. These results show that *r^y^*-yellow cultivars possess two versions of *PSY1*, one that is incomplete (*PSY1a*) and consists of a hybrid between *PSY1* and Solyc03g031870, followed by an inverted one (*PSY1b*) with the full coding sequence but with modifications in the promoter region. The inversion-duplication event explains why previous studies found two versions of *PSY1* mRNA in *r^y^* cultivars (Fray and Grierson 1993, Kang *et al*. 2014, Chattopadhyay *et al*. 2021).

**Figure 5.**
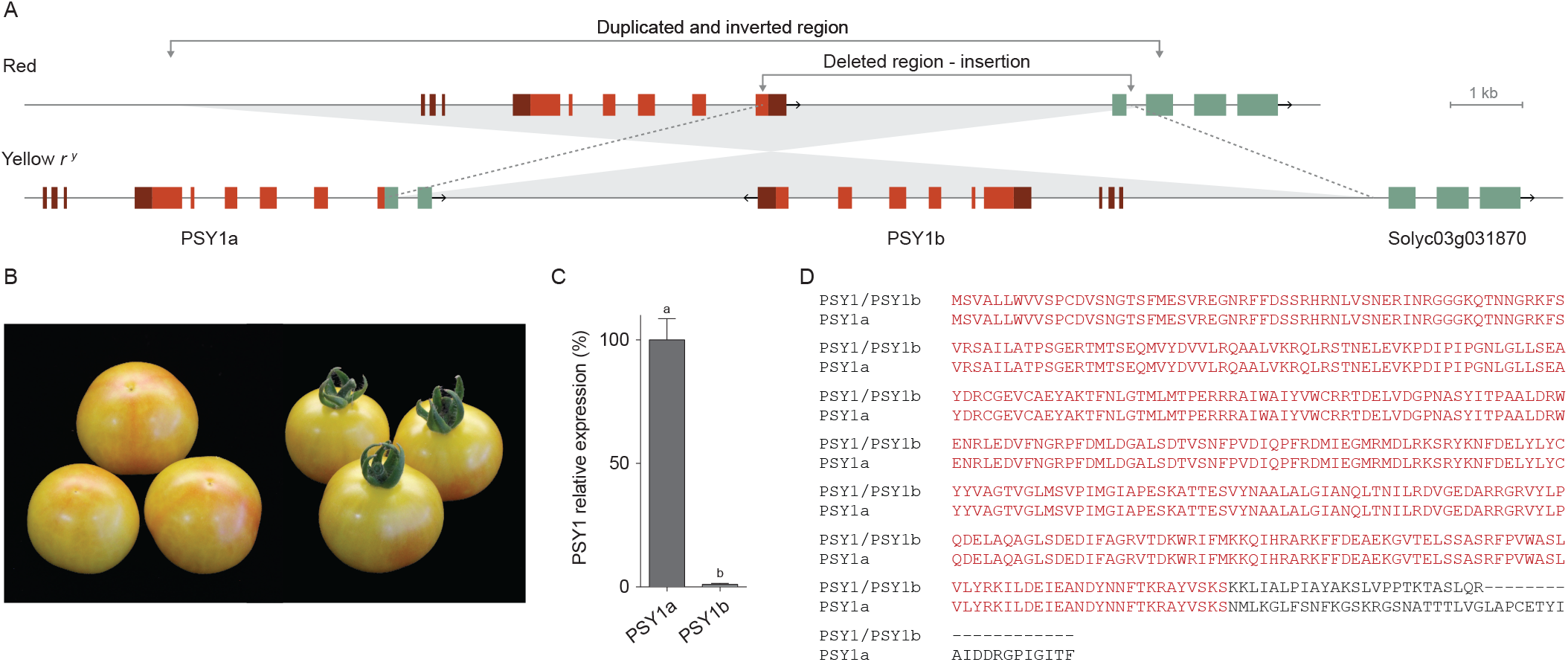
Insertion, duplication and inversion are responsible for the yellow *r^y^* phenotype in tomato cultivars. **(A)** Illustration of the genomic rearrangement associated with yellow *r^y^* cultivars. A section spanning from the promoter region of *PSY1* to the second exon of the gene located after *PSY1* (Solyc03g031870) was duplicated, inverted, then inserted in the last exon of *PSY1*. The insertion of the duplicated region led to a deletion between the last exon of *PSY1* and the first intron of Solyc03g031870. The rearrangement resulted in two copies of *PSY1*. In the first one (*PSY1*a), the end of the coding sequence was replaced by inverted parts of the gene Solyc03g031870. The end of the protein sequence of PSY1a and PSY1 is therefore different with missing amino acids and a new longer C-terminus end **(D)**. The coding sequence of the second one (*PSY1b*) is the same as the wild type *PSY1*, but the gene lacks part of the promoter region. **(C)** While *PSY1a* is highly expressed, reads specific to the end of each *PSY1* version indicate that *PSY1b* is lowly expressed indicating that the duplicated promoter region of *PSY1* is missing critical parts. **(B)** Cultivars with those two copies of *PSY1* are mostly yellow with occasional red sections near the epidermis at the blossom-end of the fruit.

RNAseq analysis shows that the transcripts from the PSY1a version of *r^y^* cultivars lack the last 76 bp of the coding sequence. These missing nucleotides are replaced by a sequence coming from the inverted gene downstream of *PSY1* (Solyc03g031870). This results in a longer coding sequence with a segment of 43 amino acids not found in the original protein sequence. The transcripts show a splicing site that starts inside Solyc03g031870 second exon and finish inside its first exon at a different position from the non-inverted sequence. The stop codon resulting from this structural variation is located inside the first exon of Solyc03g031870 (**Fig. 5**). The transcript sequence of PSY1a is identical to the cDNA sequence of the r^y^ mutant published by Fray and Grierson (1993). Yellow cultivars in the *r^y^* group have a *PSY1* gene expression reduction of around 60% compared to red cultivars. Both yellow groups show a decrease in *PSY1* expression, but the cultivars in the *r^y^* have more transcripts than the cultivars in the *r* group (**Fig. 2**). By looking at read counts for the specific end-sequence of *PSY1a* and *PSY1b* in *r^y^* cultivars, we can see that *PSY1b* expression is only 1% of the expression of *PSY1a* (**Fig. 5**). This indicate that the ~5000 bp duplicated in the promoter region is not enough to lead to a high expression. Combined with the high expression in bicolor cultivars despite a deletion at - 4858 bp, this suggest that the most important promoter regions are far from the start codon, probably starting from - 5000 bp. The weak expression of the fully functional PSY1b explains likely the slight accumulation of lycopene in parts of the fruit of *r^y^* cultivars.

### Influence of the bicolor and *r^y^* alleles of PSY1 on the fruit phenotype

Since the cultivars from the *r^y^* group can show red tissues under some conditions, crosses were made between a cultivar from the bicolor group (Granada) and a cultivar from the *r^y^* group (Azoychka). Using molecular markers, F2 homozygous plants for each allele were selected and grown for phenotyping. The plants with the *r^y^* allele were yellow with occasional red section near the blossom end, similarly to the yellow parent Azoychka. In contrast, the plants with the bicolor allele had more red tissues, especially near the epidermis, in the columella, and in the septa (**Fig. 6**). There was however variation in the intensity of the coloration in each group indicating that genes outside of the locus can influence the accumulation of lycopene for both alleles.

**Figure 6.**
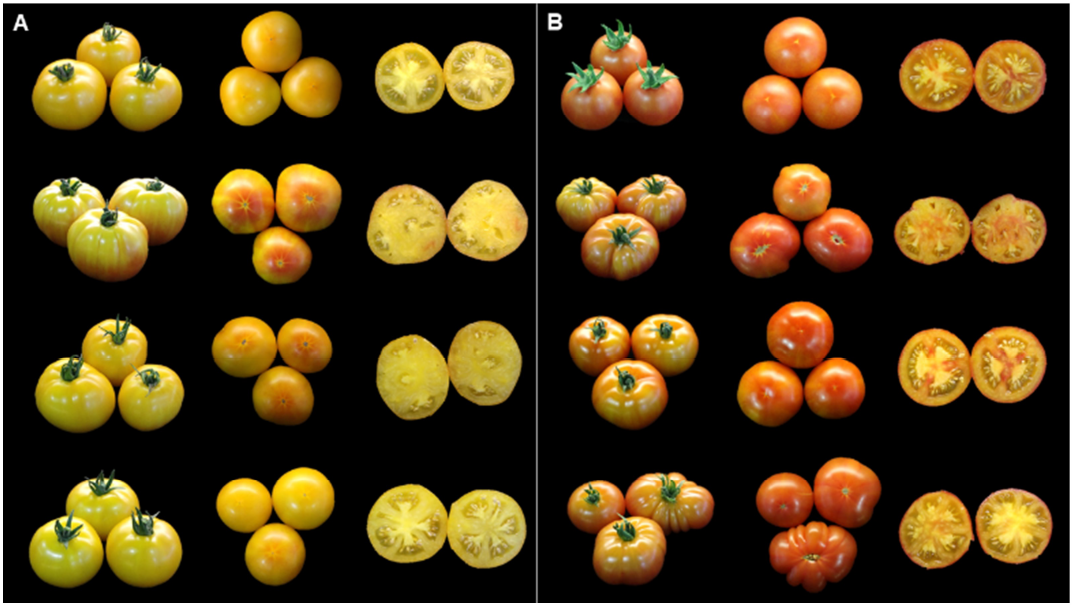
Influence of PSY1 *bicolor* and *r^y^* allele on the fruit phenotype. Yellow (**A**) and bicolor fruits (**B**) from a F2 population derived from a cross between the cultivar Azoychka (yellow *r*^y^) and the cultivar Granada (bicolor). The lines homozygous for the *r^y^* allele **(A)** are mostly yellow with occasional red tissues near the blossom end of the fruit. In contrast, the lines homozygous for the bicolor allele **(B)** show more red sections, especially near the epidermis, in the columella and in the septa.

## Discussion

Green-fruited species of the tomato clade have been useful to identify QTL involved in fruits color and to study genes of the carotenoid pathway (Liu *et al*. 2003). For example, introgressions of *S. pennellii* with orange fruits were used to identify and characterize two lycopene cyclases, LCYE (*Delta*) (Ronen *et al*. 1999) and LCYB (*Beta*) (Ronen *et al*. 2000). The same population was used to better understand the *r* locus, where the introgression of a segment of *S. pennellii* containing *PSY1* resulted in a strong decrease of carotenoids accumulation (Liu *et al*. 2003). In this study, we used introgression lines derived from the green-fruited *S. habrochaites* to fine-map a QTL resulting in bicolored fruits. The phenotype was linked to a small region on chromosome 3 where *PSY1* is located. The *S. habrochaites* introgression causes an important reduction in *PSY1* expression resulting in less accumulation of carotenoids. This reduction is likely the result of differences in the promoter of *ShPSY1* influencing transcription. Several single nucleotide polymorphisms, small deletions, and small insertions were detected in the promoter region of *ShPSY1*, disturbing many potential binding sites for transcription factors. In tomato, several ripening-related transcription factors including RIN and FUL-1 are associated with an increase in *PSY1* expression (Martel *et al*. 2011, Fujisawa *et al*. 2014, Shima *et al*. 2013). It is also possible that sequence variations between the promoter of *SlPSY1* and *ShPSY1* change the methylation pattern during the ripening. Silencing of a demethylase (*SlDML2*) in tomato results in a hypermethylation in the promoter of *PSY1* and a reduction of its expression, illustrating the importance of methylation in the control of the carotenoid pathway during the fruit ripening (Liu *et al*. 2015).

In the bicolor introgression line, the decrease of *PSY1* expression is associated with a reduction in lycopene synthesis but also a more localized accumulation of the red pigment. Interestingly, *PSY1* is not more expressed in the red sections of the fruit than in the yellow sections. This suggests either a spatial-dependent post-transcriptional regulation of PSY1 or differences in the flux inside the carotenoid pathway between sections of the fruit.

In contrast to the bicolor introgression lines derived from *S. habrochaites*, bicolor cultivars do not present a drastic reduction of *PSY1* expression. It seems unlikely that the measured reduction in *PSY1* transcripts could alone explain the phenotype. Bicolor cultivars show a structural variation in the promoter region of *PSY1* that results in a modified 5’ UTR. The transcripts of bicolor cultivars lack the first part of the 5’ UTR of red cultivars and are spliced differently resulting in a longer 5’ UTR. The sequences of 5’UTR regions are critical for gene translation efficiency by influencing, among other things, the mRNA secondary structures and the content of upstream open reading frames (Bailey-Serres 1999, Wilkie *et al*. 2003). In *Arabidopsis thaliana*, it has been demonstrated that the 5’UTR region context can make gene translation vary over 200-fold (Kim *et al*. 2014). The presence of a longer *PSY1* 5’UTR sequence in the bicolor cultivars could result in a decrease of PSY1 translation and less production of carotenoids in the fruit. In plants, most of the genes have a 5’UTR of less than 500 bp and only a small percentage have long 5’UTR of more than 1,000 bp. In *A. thaliana*, only 0.02% of genes have a 5’UTR of more than 2,000 bp (Srivastava *et al*. 2018). Given their low abundance, it is likely that longer UTR are reducing the efficiency of the translation. This can be illustrated with the two 5’UTR splicing variants for *A. thaliana PSY*. The longer variant forms an inhibitory structure that decreases the translation efficiency of *PSY* while the shorter variant allows a quick production of the protein. Analysis of truncated versions of the longer variant allowed the identification of a hairpin loop responsible for the difference between the two variants (Álvarez *et al*. 2016). With nearly 2000 nucleotides in length, the 5’ UTR of tomato bicolor cultivars is likely to form complex secondary structures that could decrease *PSY1* translation and results in the low accumulation of lycopene.

As with bicolor cultivars, *PSY1* structural variations are responsible for yellow tomatoes. In the group of yellow *r* cultivars, an insertion interrupts the first exon of *PSY1* resulting in a truncated and non-functional PSY1 protein (Fray and Grierson, 1993). The carotenoid production pathway is blocked at the first step of the pathway and the tomatoes do not accumulate carotenoids during the ripening process. This inserted sequence consists of one of the two long terminal repeats (LTR) that flank the *Rider* Ty1-copia-like retrotransposon. About 1900 copies of the Rider retrotransposon are predicted to be present in *S. lycopersicum* and some of them are still active, which means that they continue to bring major changes to its genome. For example, one *Rider* transposon is known to be responsible of the duplication and transposition of the *Sun* gene from chromosome 10 to a new locus in chromosome 7 (Jiang *et al*. 2009). A mutation in the *Rider* LTR2 induced an error during the transcription termination that usually occurs in the 3’LTR, which led to read-through transcription and the relocation of the *sun* gene along with the transposon. In this new locus, the promoter of a defensin gene disrupted by the transposition enhanced *sun* gene expression. This structural chromosomic rearrangement had an important impact on tomato shape diversity since it resulted in the apparition of oval and elongated fruits. In *PSY1* however, the sequences of the second LTR and internal coding region that are usually characteristic of retrotransposons are absent. Such solo-LTRs are not rare and constitute almost half of *Rider* retrotransposon that exist in the tomato genome (Jiang *et al*. 2009). For example, a solo-LTR is present in the *sun* locus of chromosome 7, upstream of the *sun* gene. Solo-LTRs originate by excision from the genome of one LTR and the internal coding sequence due to a recombination between the 3’ and 5’ LTRs. This special recombination is often a regulation mechanism in order to prevent insertion of too many retrotransposons and slow down genomes size expansion (King Jordan and McDonald 1999). Transposable elements are very abundant in plants genome and their insertion in gene exons often results in protein major changes or loss-of-function (Zeng and Cheng 2014, Dubin *et al*. 2018). Transposons have already been identified to cause color changes in fruits. For example, a Copia-like retrotransposon (*Tcs1*) is responsible for the purple color of blood oranges pulp (Butelli *et al*. 2012). *Tcs1* insertion near the transcription factor *Ruby* leads to activation the anthocyanin pathway in the blood oranges (Butelli *et al*. 2012). Another example is found in several cultivars of white-skinned grapes. The insertion of the transposon *Gret1* in the promoter region of the transcription factor *VvmybA1* prevents its expression and the activation of the anthocyanin pathway. In maize, kernels color is often linked to events of transposable elements that insert in genes involved in anthocyanin synthesis. One of the better-known examples is the insertion of the non-autonomous Ds transposon in an UDP-glucose flavonoid 3-O-glucosyl transferase gene (*bz* locus). The insertion results in no anthocyanin accumulation in the kernel. In the presence of the autonomous *Ac* transposon, the *Ds* transposon can move and restore the gene function causing an unstable phenotype from kernel to kernel (McClintock 1950, Fedoroff 1984).

In the *r* group of yellow cultivars, the lack of PSY1 activity comes from the insertion of a *Rider* LTR sequence, but also from an unusual alternative splicing from the end of 5’UTR exon 3 to the second PSY1 exon. In this splicing variant, the first exon is completely removed, and the transcripts don’t have the Rider solo-LTR sequence. Without the first exon, the transcripts are unlikely to result in a PSY1 functional protein. Since we didn’t detect this unusual splicing in the red, bicolor or yellow r^y^ cultivars, we can postulate that the LTR insertion is responsible for the alternative splicing. Since both splicing variants code for non-functional enzymes in the carotenoid pathway, the fruits remain yellow.

The *r^y^* yellow flesh mutation also affects PSY1 amino acid sequence. Genomic data of *PSY1 r^y^* cultivars confirmed that a chromosomic rearrangement happened, creating a complex structural variant made of two versions of *PSY1*. The inversion, duplication, and insertion created a *PSY1* version (*PSY1a*) with an altered ending. The coding sequence of the second *PSY1* (*PSY1b*) is the same as the one found in red fruits, but the gene lack some of the upstream sequences needed for a full expression. The *r^y^* cultivars have the capacity in certain conditions to accumulate low levels of lycopene at the blossom end of the fruits. This lycopene accumulation likely come from the lowly expressed PSY1a or possibly from PSY1b potentially retaining some of its activity. The latter would be consistent with the results of Kang *et al*. 2014. When expressed in bacteria, the PSY1 of r^y^ cultivars was able to successfully produce carotenoid but to a lesser extent than the wild type PSY1 (Kang *et al*. 2014). Crosses between the *r^y^* cultivar Azoychka and the bicolor cultivar Granada confirmed that the *r^y^* fruits can present red sections, but at a very low intensity compared to bicolor cultivars.

This study draws attention on the many structural variations that happened near a single gene, *PSY1*, and how those variations may lead to a diversity of colors and aromas in tomato. While bicolored tomato fruits have been described early on, the genetic basis for this unusual phenotype was still unknown. An important deletion, combined with the chromosomal rearrangement observed in *r^y^* yellow cultivars and the transposon insertion in the *r* yellow cultivars, here provides a good example of multiple phenotypes that can arise from structural variations in key steps of secondary metabolism pathways.

## Materials and Methods

### Growth of plant materials

All tomato introgression lines and cultivars were grown in University Laval, Québec. Tomato seeds were first planted in a greenhouse for germination. Plants selected were then grown in a greenhouse or in the field until they produced ripe fruits. The bicolor introgression line LA3923 is derived from a cross between *S. lycopersicum* (cultivar E6203, accession LA4024) and *S. habrochaites* (LA1777) (Monforte and Tanksley 2000). A new introgression population was then developed between *S. lycopersicum* (cultivar E6203, accession LA4024) and LA3923. The two parents and the F1 generation were grown in the field. The F2 and F3 generations of this new introgression population were respectively grown in greenhouse and in the field. F1 and F2 generations of LA3923 previously selected for possessing a fragment of *S. habrochaites* (LA1777) genome at the top of chromosome 3 (LA3923-3 and LA3923-3a lines) or at the bottom of chromosome 2 (LA3923- 2 line) were respectively grown in greenhouse and in the field. Plants in greenhouse were grown in an entirely random design. In the field, plants were placed following randomized replicated plots with three plots of three to five plants. The *S. lycopersicum* parent (LA4024) was present in all designs as a control. Bicolor cultivars (Granada, Grapefruit, Isis Candy, Mom’s, Mortgage Lifter bicolor, Mr. Stripey and Striped German), red cultivars (Ailsa Craig, Ballada, Gigantesque, Gregori’s Altai, Homer’s German Oxheart, Mexico Midget, Pruden’s Purple, Vendor and LA4024) and yellow cultivars (Barry Crazy Cherry, Butter Apple, Gallina’s Yellow, Gold Currant, Lemon Drop, Mirabelle, Poma Amoris Minora Lutea and Téton de Vénus jaune) were grown in the field over 3 summers in randomized replicated plots of two plots of five plants. The F2 population derived from the cross between Granada (bicolor) and Azoychka (yellow *r^y^*) was grown in a greenhouse.

### Lycopene analysis

Carotenoid extraction was performed as described by Anthon and Barrett 2007 in the dark on 100 mg of freeze ripe fruits ground in powder. Lycopene absorbance was then measured by a spectrophotometer (Biotek Epoch 2, www.biotek.com) at 503 nm and converted into lycopene concentration in the fruits (mg/kgfw) (Anthon and Barrett 2007).

### Volatile compounds analysis

Ripe fruits were chopped in small cubes and 100g were placed in glass tubes as described in Liscombe *et al*. 2022. During 1 hour, hydrocarbon filtered air (Agilent, www.agilent.com) arrived at one extremity of the tube, carried volatiles released by tomatoes and led them to be trapped by a divinylbenzene resin column (HayeSep Q) placed at the other extremity of the tube. Volatiles trapped on the HayeSep Q column were eluted by a solution composed of methylene chloride and the internal standard nonyl acetate. Volatile samples were then analyzed by a gas chromatograph (Agilent 6890) with a DB-5MS UI column (30m, 0.250nm diam., Agilent, www.agilent.com). Each volatile compound (Agilent, www.agilent.com) was identified by known standards and using a gas chromatograph coupled to a mass spectrometer (GC-MS, Agilent 5977B) (Agilent, www.agilent.com). For each volatile compound, one-way ANOVA followed by Tukey tests were performed in order to assess which lines or cultivars were significantly different from the others. When necessary, statistical analyses were carried out on log10 transformed data to correct for variance homogeneity.

### Genetic markers design

InDel and HRM markers were designed by comparing the genome sequences of *S. lycopersicum* (Heinz genome assembly (Fernandez-Pozo *et al*. 2015)), and *S. habrochaites* (cv. LYC4) (SAMEA3138934, Tomato Genome Consortium 2012). Homologous sequences were aligned with MAFFT program (Katoh and Standley 2013) to find an insertion or a deletion (InDel marker), or a SNP (HRM marker). The mutation had to be flanked by conserved sequences between the two species to ensure proper amplification by PCR. The primers were designed with Primer3 (Rozen and Skaletsky 2000). For the F2 population derived from Granada x Azoychka crosses, an A to G substitution mutation presents in *PSY1* intron 4 of *r* yellow cultivars was used to predict the F2 segregation ratio with genetic markers.

### Quantitative PCR

Ripe fruits of LA4024, LA3923-3a and LA3923-2 lines were frozen in liquid nitrogen and store at -80°C. Total RNA was extracted following the EZ-10 Spin Column Plant RNA Miniprep Kit protocol (Biobasic, www.biobasic.com). Possible genomic DNA contamination was removed by DNAse treatment step (Qiagen, www.qiagen.com) before purifying total RNA with a second run of the EZ-10 Spin Column Plant RNA Miniprep Kit protocol. Quantification of total RNA was evaluated on NanoDrop (ThermoFisher, www.thermofisher.com). Quantitative PCR (qPCR) was executed with the qScript One-Step qRT-PCR Kit (QuantaBio, www.quantabio.com) from 100ng of total RNA. The primers were designed for the amplification of *PSY1* mRNA. To amplify mRNA only and not genomic DNA, the forward primer was designed to begin at the end of exon 4 and finish at the beginning of exon 5, while the reverse primer was located at the beginning of exon 6. The quantification of *PSY1* transcripts was determined with a standard curve. Expression levels have been compared with Tukey test. Significant differences are indicated with letters (*p*<0.05).

### Transcriptome sequencing and analysis

Total RNA was extracted from ripe fruits of LA4024, LA3923-3a, red, bicolor, and yellow cultivars with the same protocol as for qPCR. The RNA-Seq libraries were prepared following the Zhong *et al*. 2011 protocol that has been modified and adapted for paired-end 125 pb RNA-Seq. The libraries were sent to McGill University sequencing platform (www.gqinnovationcenter.com) to be analyzed on a Bioanalyzer (Agilent, www.agilent.com) and sequenced paired-end 125 bp with HiSeq 4000 Illumina technology (Illumina, www.illumina.com). The quality of the sequencing data was then evaluated with FastQC program. The bad qualities reads and Illumina adapters were removed with Trimmomatic (Bolger *et al*. 2014). The reads were then aligned to the *Solanum lycopersicum* reference genome (version SL3.0) available at Sol Genomics Network (Fernandez-Pozo *et al*. 2015) or the homemade *r^y^*-yellow reference genome (see “Genomic analysis section”) with Hisat2 (Kim *et al*. 2015). Gene expression (FPKM) was assessed with Stringtie (Pertea *et al*. 2015). The packages edgeR (Robinson *et al*. 2010), DESeq2 (Love *et al*. 2014) and Limma-voom (Ritchie *et al*. 2015) from the Bioconductor repertory of R (Bionconductor, www.bioconductor.org) were then used to perform the differential gene expression analysis. *PSY1* expression levels have been compared with Tukey test. SNPs calling in LA4024 and LA3923-3a lines was performed with SAMtools and bcftools (Danecek et al. 2021), and SNP density was evaluated with Sniplay (Dereeper *et al*. 2011). The program regtools was used in order to extract the splicing variants that can be present in each samples, and visualization was performed thanks to the IGV software (Robinson *et al*. 2011). Yellow cultivars PSY1 sequences were compared with *r* (X67143.1) and *r^y^* (X67144.1) mutant sequences available on NCBI.

### Genomic analysis

SRA data of tomato bicolor and yellow *r^y^* cultivars from project PRJNA353161 (Tieman *et al*. 2017) and PRJNA423252 (University of Georgia) were analyzed to study their respective mutations. As for RNA-seq samples, data were trimmed to remove bad quality reads and Illumina adapters. Trimmed data were then aligned to the *Solanum lycopersicum* reference genome (SL3.0) with Hisat2 program and the alignment result was visualized with IGV. The yellow *r^y^* data were also used to carry out a *de novo* genome assembly of the tomato *r^y^* genome with ABySS program (Simpson *et al*. 2009). This assembly helped to create a *r^y^*-yellow reference genome built from *S. lycopersicum* reference genome (SL3.0).

## Acknowledgment

This work was supported by funding from Fonds de recherche du Québec – Nature et technologies, Genome Quebec and Genome Canada. Computations were made on the supercomputer Mammouth-Mp2 from Sherbrooke University, managed by Calcul Québec and Compute Canada. The operation of this supercomputer is funded by the Canada Foundation for Innovation (CFI), the ministère de l’Économie, de la science et de l’innovation du Québec (MESI) and the Fonds de recherche du Québec - Nature et technologies (FRQ-NT).

