## Supplementary Fig. 1 for "Structural variations in the phytoene synthase 1 gene affect carotenoid accumulation in tomato fruits and result in bicolor and yellow phenotypes"

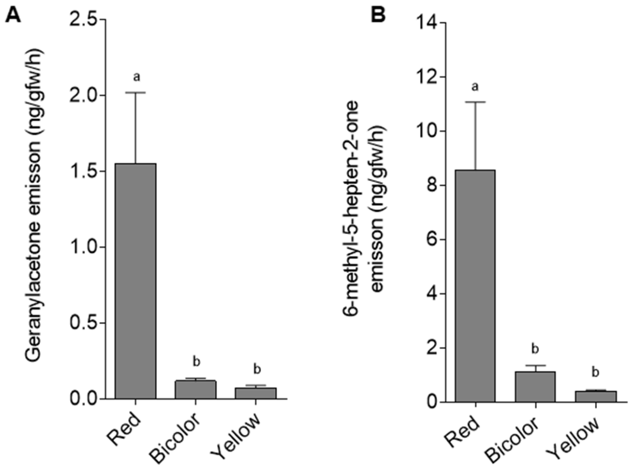


**Supplementary Figure 1. Apocarotenoid volatiles emission from fruits of red, bicolor, and yellow cultivars.** Emission of the apocarotenoids geranylacetone **(A)** and 6-methyl-5-hepten-2-one **(B)** (±SE, $p<0.05$, n=4).
